# XPC loss-of-function triggers melanomagenesis through *CDKN2A* downregulation

**DOI:** 10.1101/2025.04.03.646637

**Authors:** Vipin Shankar Chelakkot, Kiara Thomas, Todd Romigh, Alexis Bernat, Reina Samuel, Nihit Goli, Pooja Rambhia, Sairekha Ravichandran, Qi-En Wang, Kevin D. Cooper, Pauline Funchain, Ying Ni, Joshua Arbesman

**Author notes:** **Corresponding Author:** Joshua Arbesman.

## Abstract

We identified a novel XPC variant, c.2420+5G>A (XPCvar), in siblings with multiple melanomas, inherited alongside c.779+1G>T, which results in an absent or disrupted protein. However, they did not exhibit significantly higher nucleotide excision repair (NER) deficits compared to their unaffected parents, suggesting an NER-independent tumor suppressor function for XPC. XPC knockdown increased cell proliferation and tumorigenicity in vitro without affecting NER. Single-cell RNA sequencing revealed lower Cdkn2a in Xpcvar/var mouse melanocytes. Additionally, XPC knockdown downregulated CDKN2A in vitro. Furthermore, patient fibroblasts showed decreased p16INK4a, which was rescued by XPC overexpression. ChIP-PCR and luciferase assays confirmed XPC binding to the CDKN2A promoter, initiating transcription. Premature stop codon read-through, with gentamicin, restored XPC and p16INK4A in patient fibroblasts. These suggest that XPC regulates CDKN2A and XPC loss might promote melanomagenesis by downregulating CDKN2A, independent of its NER function. We provide preclinical evidence for potential preventive and/or therapeutic strategies for similarly affected individuals.

**STATEMENT OF SIGNIFICANCE:** Characterizing a novel *XPC* variant identified in patients with melanoma revealed a non-canonical role for XPC in regulating *CDKN2A* transcription. XPC binding to the *CDKN2A* promoter is crucial for *CDKN2A* expression and might suppress melanomagenesis. This expands our understanding of melanomagenesis and suggests potential therapies targeting XPC-mediated regulation of *CDKN2A*.

## INTRODUCTION

Melanoma, a highly aggressive form of skin cancer, presents a significant health challenge due to its rapid progression and high mortality rates(1,2). While germline genetics drive melanoma susceptibility(3,4), the cellular mechanisms underlying this predisposition remain elusive(4–6). Characterizing these mechanisms enables the development of targeted therapies and preventive strategies tailored for patients carrying specific germline gene variants, as demonstrated by a previous study using Poly-ADP ribose polymerase (PARP) inhibitors(7). Furthermore, elucidating these germline mechanisms would facilitate early melanoma detection and personalized treatment, improving patient outcomes.

We recruited two siblings diagnosed with multiple melanomas at a relatively young age, along with their immediate family, to characterize the cellular mechanisms contributing to melanomagenesis in very high-risk individuals. Whole-genome sequencing identified a novel variant of Xeroderma pigmentosum, complementation group C (*XPC*), a gene traditionally associated with nucleotide excision repair (NER)(8,9) inherited alongside an *XPC* loss-of-function variant. However, the patients did not exhibit significant DNA repair deficits compared to their unaffected parents, leading us to hypothesize that *XPC* has an additional function relevant to melanoma development beyond its role in DNA repair. Our results demonstrated that XPC regulates *CDKN2A* transcription, independent of DNA repair, revealing a novel tumor suppressor function in melanocytes. This patient-driven study revealed a global gene regulatory mechanism that could illuminate broader aspects of gene regulation and cancer pathogenesis.

## RESULTS

### A novel XPC variant was identified from a patient family with multiple melanomas

We encountered two siblings who presented with multiple (>10) melanomas at a relatively young age, accompanied by atypical/dysplastic nevi (Fig.1A). Their first melanomas were diagnosed at ages 17 and 21, with several subsequent melanomas and dysplastic nevi identified. Both siblings exhibited a high frequency of lentigines in sun-exposed areas but did not have any additional unique skin findings. The sister developed one basal cell carcinoma, but no other keratinocyte cancers (basal cell carcinoma or squamous cell carcinoma) were found in either sibling. These findings raised suspicion of Familial Atypical Multiple Mole Melanoma Syndrome, a hereditary cancer syndrome characterized by multiple melanomas, numerous atypical nevi, and a strong family history of melanoma. Regardless of the specific syndrome, this family clearly exhibited a significant genetic predisposition to melanoma. Accordingly, the patients and their immediate family members were enrolled in our study after procuring signed informed consent. Blood was collected for whole-exome and/or whole-genome sequencing, and punch biopsies were obtained of normal, unaffected, non-sun-exposed skin to obtain fibroblasts for functional studies.

**Figure 1.**
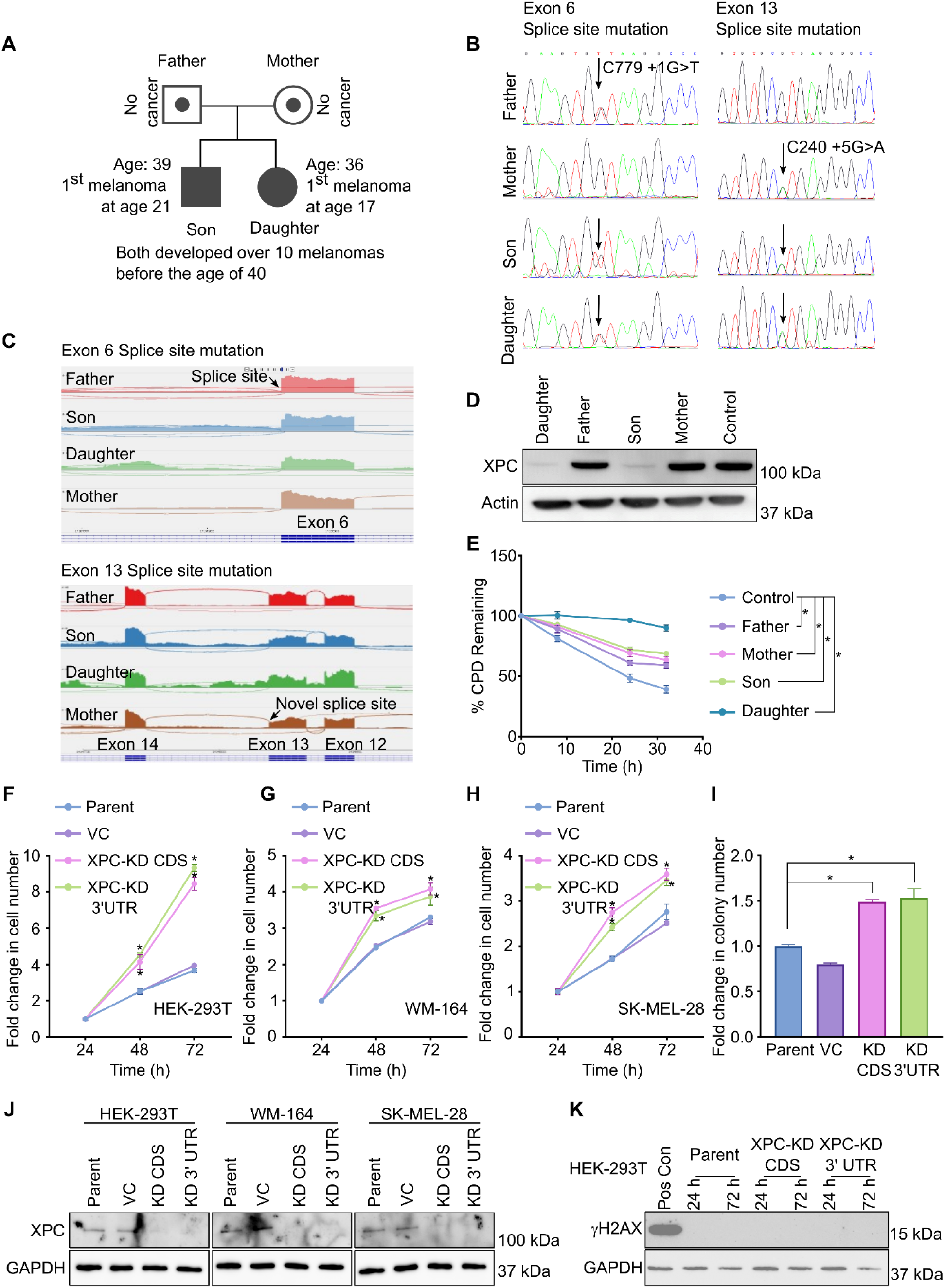
A novel *XPC* variant, c240+5G>A, was identified from two sibling patients who presented with multiple melanomas before the age of 40. (**A**) Melanoma inheritance pattern in the patient family. Two children (a son and a daughter) developed multiple melanomas before the age of 40, while the parents have not developed any melanomas. (**B**) Whole-genome sequencing identified two XPC variants in the patients, c.2420 +5G>A inherited from the mother and c.779+1G>T inherited from the father. (**C**) Sashimi plot analysis of the possible alternative splicing products of XPC in the patients and their parents. Zoomed-out sashimi plots are shown in Fig.S1A. (**D**) Representative western blot for XPC in fibroblasts derived from the patients and their parents. Human dermal fibroblasts (HDFa) were used as the controls, and actin was used as the loading control. (**E**) Percentage cyclobutane pyrimidine dimers (CPD) retained in the fibroblasts derived from the patients and their parents and normal human skin fibroblasts (OSU-2, Control) at different time points after ultraviolet-C (UVC) irradiation. Data expressed as mean ± SEM of 3 independent experiments. *p<0.05, determined using two-way ANOVA with Tukey’s multiple comparisons. Representative gel image is shown in Fig.S1B (**F-H**) Fold change in cell numbers of (**F**) HEK-293T, (**G**) WM-164, and (**H**) SK-MEL-28 parent, vector control (VC), or XPC knockdown (knockdown of the coding sequence (XPC-KD CDS) or the 3′ UTR (XPC-KD 3′ UTR) cells with time. Data expressed as mean ± SEM of 3 independent experiments. *p<0.05, determined using two-way ANOVA with Tukey’s multiple comparisons. (**I**) Fold change in the number of soft agar colonies formed by parent, VC, XPC-KD CDS, and XPC-KD 3′UTR HEK-293T cells at 3 weeks after plating on soft agar plates. Data expressed as mean ± SEM of 3 independent experiments. *p<0.05, determined using one-way ANOVA with Tukey’s multiple comparisons. (**J**) Representative western blot for XPC on the parent, VC, XPC-KD CDS, and XPC-KD 3′UTR HEK-293T, WM-164, and SK-MEL-28 cells. GAPDH was used as the loading control. (**K**) Representative western blot for γH2AX, a marker of DNA damage, on HEK-293T parent, XPC-KD CDS, and XPC-KD 3′UTR cells at two different time points in the experimental window.

Whole-genome sequencing identified the patients as compound heterozygous for *XPC* harboring two *XPC* variants, c.2420 +5G>A, inherited from the mother, and c.779+1G>T, inherited from the father (Fig.1B). A variant at the same site as the c.779+1G>T variant has been reported to affect a donor splice site in intron 6 of *XPC* and is expected to disrupt RNA splicing, resulting in loss of XPC function, and has been classified as likely pathogenic or pathogenic(10–13). The c.2420 +5G>A variant has not been formally reported before and is a novel variant that we identified in the patients.

Whole transcriptomes from tumor tissues were analyzed for *XPC* isoforms and presented as Sashimi plots to investigate the impact of these variants on RNA splicing and to detect aberrant splicing events (Fig.1C,S1A,B). Analysis of the c.2420+5G>A variant carrier samples revealed disrupted splicing, leading to the introduction of a premature termination codon (PTC) downstream of exon 13, which would result in a truncated, likely non-functional protein. Additionally, the sashimi plots indicated potential retention of the whole or parts of intron 13, consistent with defective splicing at this site. Western blot analysis of fibroblasts from the patients and their parents showed XPC levels similar to that in control healthy fibroblasts in the parents and markedly lower XPC levels in the patients. Interestingly, XPC bands with altered molecular weights were not observed in the patient fibroblasts (Fig.1D). This suggested that the truncated protein might lack stability and could be degraded by the cells. Nevertheless, the RNA was likely not degraded secondary to nonsense-mediated decay, given the position of the PTC in the mRNA. Additionally, the faint XPC bands observed in the patient fibroblasts indicated that the impact of the splice site variant may not be complete and that there is low-level production of XPC. Whole-exome and whole-genome sequencing did not identify any additional significant variants in the patients, suggesting that the low XPC levels are likely relevant to melanomagenesis in these patients.

### The novel XPC variant is sufficient for basal DNA repair function

XPC loss-of-function is traditionally associated with reduced NER efficiency, which is thought to result in an accumulation of UV-induced mutations and the subsequent development of cancers(14–16). We used a cyclobutane pyrimidine dimer (CPD) assay to estimate the DNA repair capacity of fibroblasts from patients and their parents to determine whether low XPC levels affected their NER capacity (Fig.1E,S1C). We compared the rate at which fibroblasts from the patients and their parents repaired UV-induced CPDs with that of normal human skin fibroblasts, OSU-2. The control fibroblasts showed a 60% decrease in UV-induced CPDs at 32 h after UV exposure. However, the fibroblasts from both the patients and their parents exhibited a significant accumulation of UV-induced CPDs at 32 h post-UV exposure. The patient fibroblasts retained approximately 95% and 70% of the CPDs, while the fibroblasts from the parents retained approximately 65% of the CPDs at 32 h post-UV exposure and showed no significant difference compared to the patient-derived fibroblasts (Fig.1E,S1C). This suggested that all family members had some degree of NER deficiency compared to control skin fibroblasts. Furthermore, the patients did not show a significantly higher NER deficit compared to their parents. Surprisingly, the NER deficit in the parents did not lead to melanomagenesis, while the patients developed multiple melanomas before turning 40 years old. Additionally, the patients did not exhibit additional phenotypic changes typically associated with NER deficiency, such as the development of multiple basal and squamous cell carcinomas. These indicated an NER-independent pathway for melanomagenesis in the patients.

### XPC loss might contribute to melanomagenesis, independent of its NER function

To evaluate whether XPC downregulation alone would contribute to carcinogenesis, we knocked down endogenous XPC in the normal cell line HEK-293T and two melanoma cell lines, WM-164 and SK-MEL-28. Knocking down XPC significantly increased cell proliferation in all three cell lines and increased in vitro tumorigenicity in HEK-293T cells (Fig.1F-J). To determine whether the cells accumulated DNA damage during the experiment period, we collected cells at different time points within the experiment window and determined the expression of γH2AX, a marker for DNA damage, by western blotting (Fig.1K). The cells did not express γH2AX at any of the time points within the experiment window, suggesting that the increase in proliferation and tumorigenicity was not a result of accumulated DNA damage.

### Xpc^var/var^ mice did not express Cdkn2a in their tail skin melanocytes

To characterize the specific effect of the novel *XPC* variant identified from the patient family, we developed a mouse model carrying the *Xpc* splicing variant (*Xpc*^var/var^), representative of the human *XPC* variant identified in the patient family (Fig.S2A). Although we detected an altered *Xpc* mRNA transcript in different mouse tissues by PCR, we were unable to identify it at the protein level due to the lack of a reproducible mouse *Xpc*-variant-specific antibody. Additionally, PCR evaluation of the *Xpc* gene indicated that the mice expressed the wild-type (WT) form along with the intron 13-retaining form (Fig.S2B). The *Xpc^var/var^ m*ice did not show any phenotypic changes and did not develop any adverse conditions, even with aging.

To determine whether the altered *Xpc* had any skin cell-specific effects, we performed a single-cell RNA-seq analysis of mouse tail skin (Fig.2A). We chose tail skin because its melanocyte arrangement resembles that of human skin. Cell clusters were identified using the shared nearest neighbor (SNN)(17) modularity optimization-based clustering algorithm in Seurat, as described previously(18). SNN-based clustering at an SNN resolution of 0.9 identified 26 unique cell clusters. Single-cell RNA-seq analysis revealed a similar composition and distribution of skin cells in the WT and Xpcvar/var mice (Fig.2A). Cell-type annotations were determined using the Single-Cell Scoring Annotations (SCSA), which integrates differentially expressed genes and confidence levels of cell markers from multiple databases. Importantly, SCSA effectively annotated the 26 cell clusters, providing a comprehensive representation of the cell composition typically observed in mouse tail skin.

**Figure 2.**
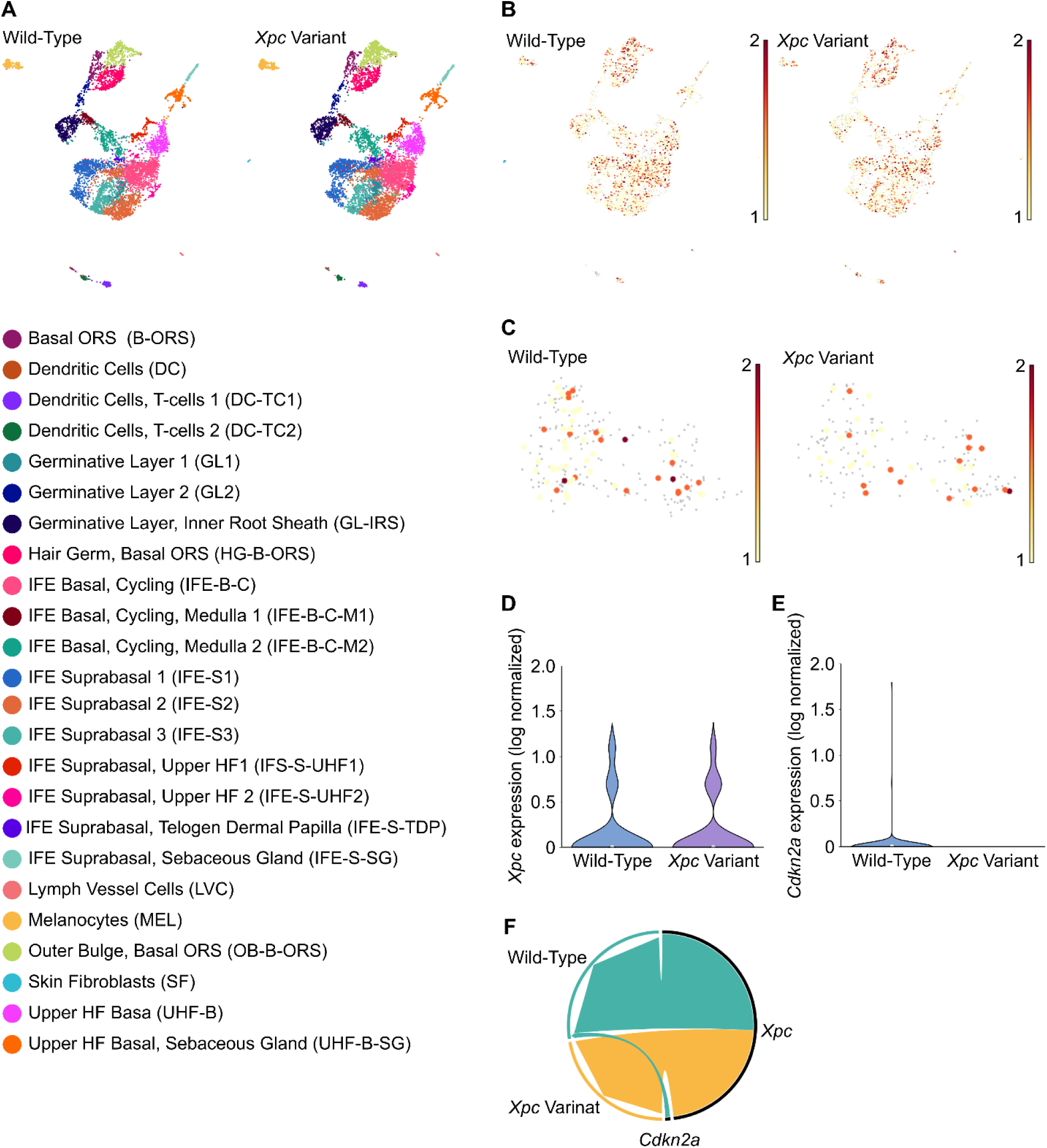
Xpc^var/var^ mice did not express *Cdkn2a* in their tail skin melanocytes. (**A**) UMAP displays unsupervised clustering of all cells, identified using the shared nearest neighbor (SNN) modularity optimization-based clustering algorithm in Seurat, in the wild-type and Xpc^var/var^ mice. The cell-type annotation of the clusters is based on established marker genes, as determined by the SCSA analysis, with clusters listed alphabetically. The number in the cluster name (e.g., T-cells 1 and T-cells 2) shows that the clusters were defined by different gene sets. The cell-type annotation of the clusters is based on established marker genes, as determined by the SCSA analysis, with clusters listed alphabetically. (**B**) UMAP displays *Xpc* expression levels in the different clusters of the wild-type and Xpc^var/var^ mice. (**C**) UMAP displays Xpc expression levels in melanocytes of the wild-type and Xpc^var/var^ mice. (**D**) Violin plot showing *Xpc* expression within melanocytes of wild-type and Xpc^var/var^ mice. (**E**) Violin plot showing *Cdkn2a* expression within melanocytes of wild-type and Xpc^var/var^ mice. (**F**) Circos plot showing the relationship between *Xpc* and *Cdkn2a* expression in the melanocytes of wild-type and Xpc^var/var^ mice.

Analysis of gene expression levels in different clusters did not identify any significant differences in *Xpc* expression in any of the skin cell clusters between WT and *Xpc^var/var^* mice (Fig.2B,S3A), including the melanocytes (Fig.2C,2D). Analysis of differentially expressed genes in melanocytes from WT and *Xpc^var/var^* mice revealed that the *Xpc^var/var^ mice* did not express *Cdkn2a* in their melanocytes (Fig.2E), whereas the WT mouse had a distinct *Cdkn2a*-expressing melanocyte cluster (Fig.2F). Interestingly, several clusters did not express *Cdkn2a* (Fig.S3B). Three clusters (Dendritic cells-T-cell 2, Germinative layer-Inner root sheath, and Hair germ-Basal ORS) exhibited comparable *Cdkn2a* expression between WT and Xpc^var/var^ mice. In contrast, six clusters (Germinative layer 1, Germinative layer 2, IFE Basal-Cycling-Medulla 1, IFE Suprabasal 2, Melanocytes, and Upper HF basal Sebaceous gland) showed significantly higher *Cdkn2a* levels in WT mice, and two clusters (IFE Suprabasal-Sebaceous gland and IFE Suprabasal Upper HF 1) exhibited higher *Cdkn2a* expression in Xpc^var/var^ mice.

### XPC regulates CDKN2A expression

*CDKN2A* is a tumor suppressor gene crucial in melanomagenesis, both as a germline risk factor and through somatic events(19,20). Dysregulation of *CDKN2A*, often through genetic alterations or epigenetic modifications, contributes significantly to melanoma development and progression(19,21–24). However, the mechanisms governing *CDKN2A* expression remain largely unknown. Our single-cell RNA-seq analysis identified that the *Xpc^var/var^* mice did not express *Cdkn2a* in their melanocytes (Fig.2E,2F). Therefore, we evaluated the effect of XPC on *CDKN2A* expression through in vitro loss-of-function and gain-of-function studies in HEK-293, WM-164, and SK-MEL-28 cells (Fig.3A-D). WM-164 and SK-MEL-28 are melanoma cell lines, and HEK-293 cells were used as they represent normal cells with no oncogenic transformation(25).

**Figure 3.**
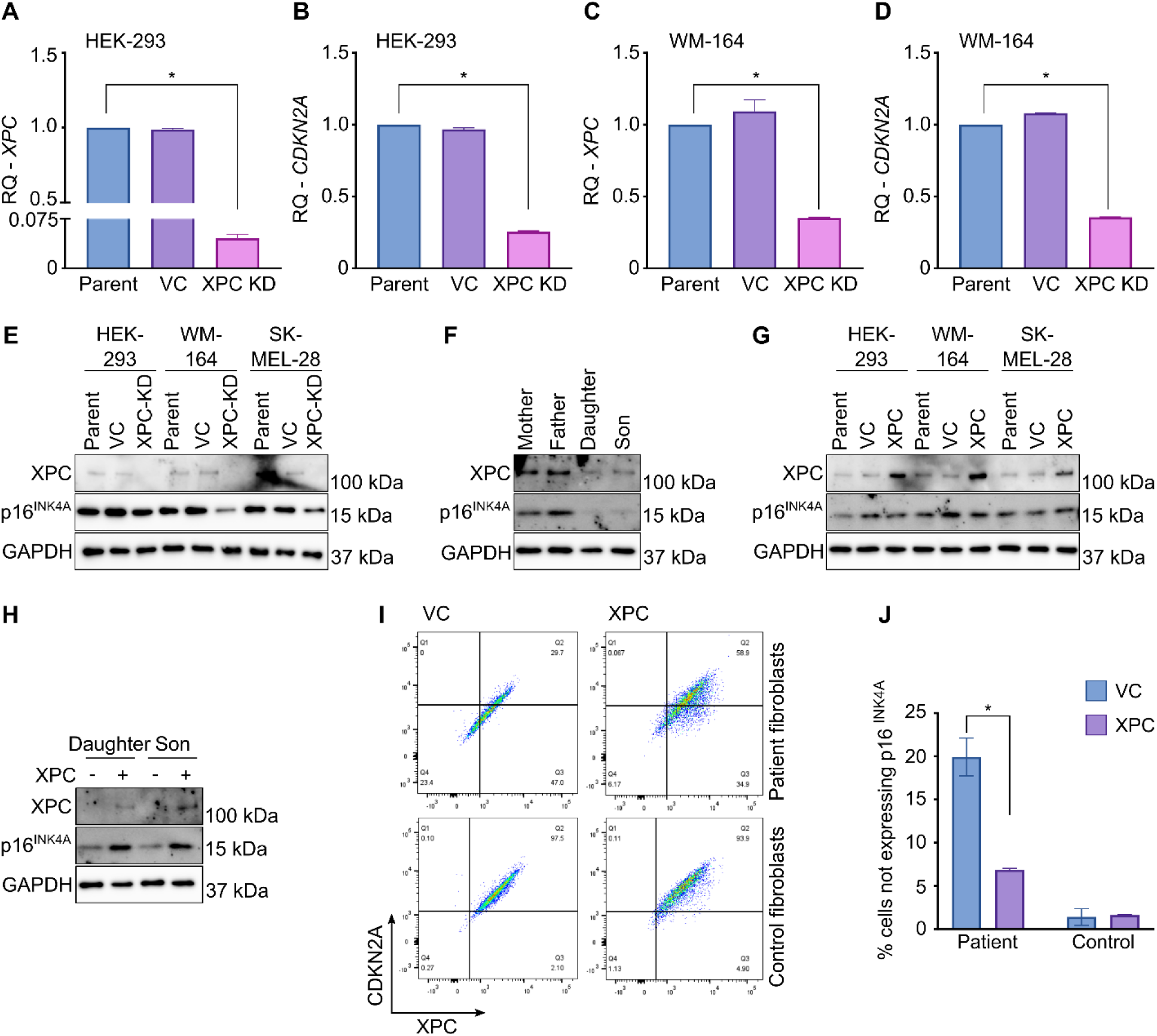
XPC regulates *CDKN2A* expression. (**A-D)**. qRT-PCR analysis for *XPC* (A, C) and *CDKN2A* (B, D) expression in HEK-293 (A, B) and WM-164 (C, D) parent, vector control (VC), and XPC knockdown (XPC-KD) cells. The expression levels were normalized to GAPDH and presented as mean ± SEM of the relative quantification (RQ) values from 3 independent experiments. *p<0.05, determined using one-way ANOVA with Tukey’s multiple comparisons. (**E**) Representative western blot for XPC and p16^INK4A^ on the parent, VC, and XPC knockdown (XPC-KD) HEK-293, WM-164, and SK-MEL-28 cells. GAPDH was used as the loading control. (**F**) Representative western blot for XPC and p16^INK4A^ fibroblasts derived from the patients and their parents. GAPDH was used as the loading control. (**G**) Representative western blot for XPC and p16^INK4A^ on the parent, VC, and XPC overexpressing (XPC) HEK-293, WM-164, and SK-MEL-28 cells. GAPDH was used as the loading control. (**H**) Representative western blot for XPC and p16^INK4A^ on the patient-derived fibroblasts transfected with either the VC or XPC overexpression plasmid. GAPDH was used as the loading control. (**I**) Representative dot plots of XPC and CDKN2A expression in control (HDFa) and patient-derived fibroblasts transfected with either the VC or XPC overexpression plasmid determined using flow cytometry. (**J**) Percentage cells not expressing p16^INK4A^ in the patient and control (HDFa) and patient-derived fibroblasts transfected with either the VC or XPC overexpression plasmid, determined by flow cytometry. Data expressed as mean ± SEM of 3 independent experiments. *p<0.05, determined using Student’s *t*-test.

Knocking down XPC downregulated *CDKN2A* mRNA in HEK-293 and WM-164 cells (Fig.3A-D) and p16^INK4A^ in WM-164 and SK-MEL-28 melanoma cells (Fig.3E). However, no p16^INK4A^ downregulation was observed in HEK-293 cells (Fig.3E). Similarly, patient fibroblasts showed lower p16^INK4A^ expression compared to fibroblasts from the parents (Fig.3F). However, overexpressing XPC in HEK-293, WM-164, and SK-MEL-28 cells did not upregulate p16^INK4A^ expression (Fig.3G). On the other hand, overexpressing XPC in patient fibroblasts with low XPC rescued p16^INK4A^ expression (Fig.3H). Additionally, flow cytometry showed that overexpressing XPC in patient fibroblasts significantly decreased the percentage of cells not expressing CDKN2A, implying an increase in the percentage of CDKN2A-expressing cells (Fig.3I-J). Interestingly, a similar change was not observed in the control fibroblasts, HDFa, where XPC upregulation did not significantly affect CDKN2A expression (Fig.3I-J). These results suggested that XPC modulated the expression of *CDKN2A* in patient fibroblasts and, to a lesser extent, in melanoma cells but not in HEK-293 or healthy skin fibroblasts.

### XPC binds to the CDKN2A promoter and is required for CDKN2A expression

XPC, beyond its well-established role in DNA repair, has been shown to function as a cofactor for RNA polymerase, suggesting a role in transcription initiation(26–28). Previous studies have demonstrated XPC’s involvement in gene regulation; however, a comprehensive list of all genes regulated by XPC is currently unavailable, and the specific DNA sequence or motif recognized by XPC for its transcriptional activity remains unidentified(26,29). Since XPC loss and gain affected *CDKN2A* mRNA and protein levels in vitro, we explored whether XPC binds to the *CDKN2A* promoter and is necessary for *CDKN2A* expression. To determine whether XPC binds to the *CDKN2A* promoter, we first pulled down XPC-bound DNA fragments and used qPCR to determine whether *CDKN2A* promoter regions could be amplified from the pull-down. The *CDKN2A* gene encodes two distinct tumor suppressor proteins, p16^INK4A^ and p14^ARF^, by using alternative promoters(30) (Fig.4A). Therefore, we used two sets of primers to target both regions (Primers 1 and 2 targeted Promoter 1, and Primers 3 and 4 targeted Promoter 2 (Fig.4A)) for the qPCR assay. Both promoter regions were amplified from the XPC-pull-down but not from the GFP-pull-down, which served as the negative control (Fig. 4B). This suggested that XPC binds to both promoter regions of *CDKN2A*.

**Figure 4.**
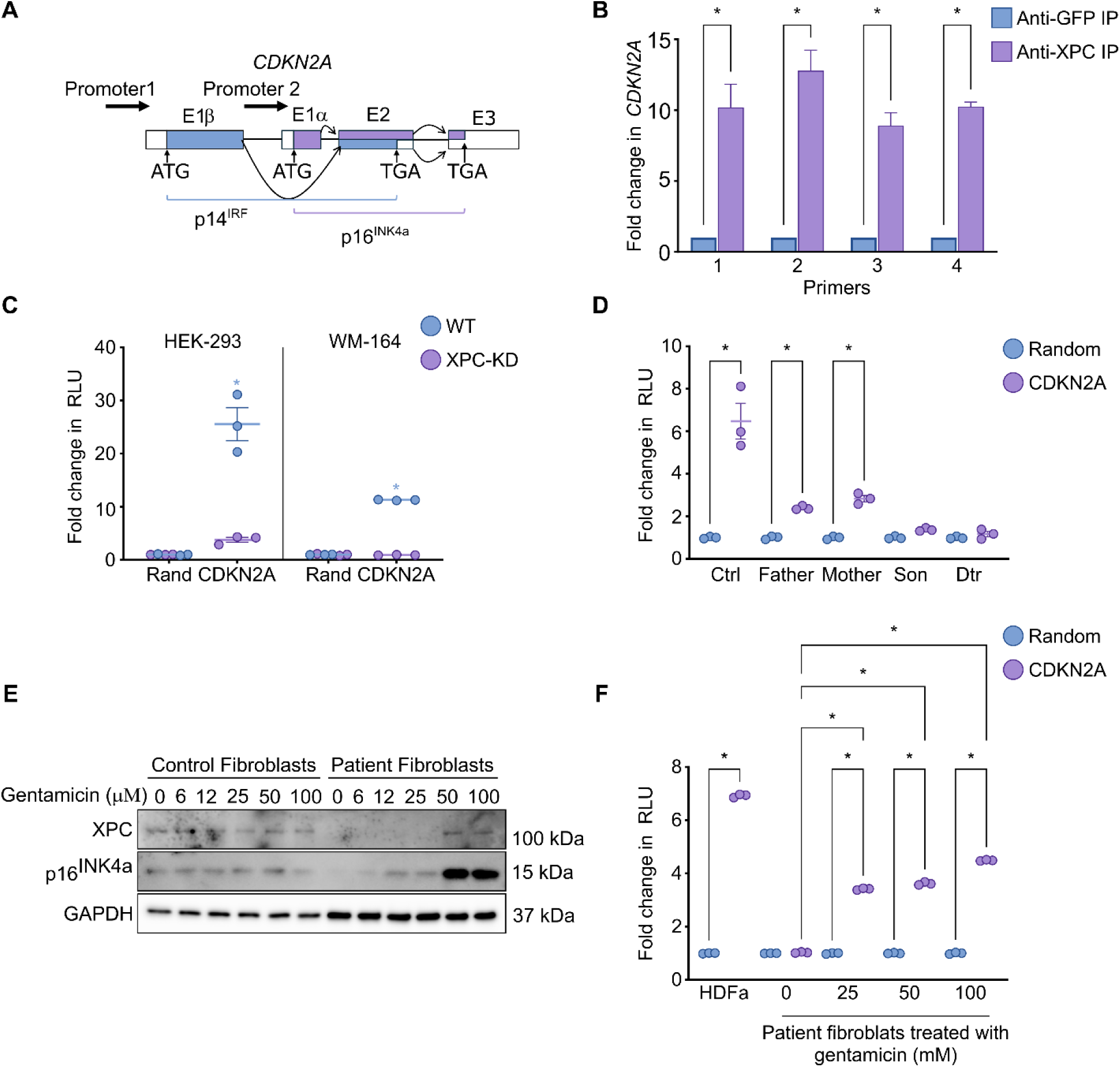
XPC binds to the *CDKN2A* promoter and is required for *CDKN2A* expression. (**A**) Schematic showing the gene structure of *CDKN2A* and the promotor regions. (**B**) ChIP fold enrichment of DNA fragments around the *CDKN2A* promoter regions by ChIP-qPCR. Four primer sets were used: Primer sets 1 and 2 targeted Promotor 1 and Prime sets 3 and 4 targeted Promotor 2 of the *CDKN2A* gene. Data expressed as mean ± SEM of 3 independent experiments. *p<0.05, determined using Student’s *t*-test. (**C**) Fold change in relative luminescence units (RLU) in wild-type (WT) and XPC-knock-down (XPC-KD) HEK-293 and WM-164 cells transiently transfected with a luciferase construct with the Promotor 2 region of *CDKN2A* or a random negative control (Rand) 48 h after transfection. Data from 3 independent experiments are shown. Error bar indicates SEM. *p<0.05, determined using two-way ANOVA with Tukey’s multiple comparisons. (**D**) Fold change in RLU in control fibroblasts and fibroblasts derived from the patients and their parents transiently transfected with a luciferase construct with the Promotor 2 region of *CDKN2A* or a random negative control (Rand) 48 h after transfection. Data from 3 independent experiments are shown. Error bar indicates SEM. *p<0.05, determined using two-way ANOVA with Tukey’s multiple comparisons. (**E**) Representative western blot for XPC and p16^INK4A^ in control fibroblasts (HDFa) and patient (Son)-derived fibroblasts treated with or without different concentrations of gentamicin. GAPDH was used as the loading control. (**F**) Fold change in RLU in control fibroblasts and patient (Son)-derived fibroblasts treated with different concentrations of gentamicin and transiently transfected with a luciferase construct with the Promotor 2 region of *CDKN2A* or a random negative control (Rand) 48 h after transfection. Data from 3 independent experiments are shown. Error bar indicates SEM. *p<0.05, determined using two-way ANOVA with Tukey’s multiple comparisons.

Subsequently, we determined whether XPC binding to the promoter sequence initiates transcription using a luciferase assay. A luciferase construct with an upstream CDKN2A promoter (Promoter 2 in Fig.4A) or a random negative control promoter was transfected into WT or stable XPC-knock down HEK-293 and WM-164 cells. The XPC-knockdown HEK-293 and WM-164 cells showed significantly lower luciferase signals compared to parental lines, while no luciferase signal was detected in the cells transfected with the negative control construct (Fig.4C), suggesting that XPC is essential for the successful transcription and translation of genes downstream of the *CDKN2A* promoter. Further, control fibroblasts and fibroblasts from parents transfected with the CDKN2A-luciferase construct produced detectable levels of luciferase, while patient fibroblasts (with low XPC levels) transfected with the construct failed to produce detectable luciferase signals (Fig.4D). These results confirmed that XPC binds to the *CDKN2A* promoter region and is required for the initiation of transcription of both *CDKN2A* proteins.

Since the XPC variant identified from the patients has a PTC that might have resulted in either an absent or truncated protein, we next attempted to bypass the PTC using gentamicin for potential clinical translation. Gentamicin, an aminoglycoside antibiotic, can induce PTC read-through(31,32). This occurs when the ribosome misinterprets a stop codon as an amino acid codon, allowing translation to proceed beyond the premature stop site(32). Treating patient fibroblasts with gentamicin restored XPC and p16^INK4A^, as determined by western blotting (Fig.4E), in a dose-dependent manner. However, gentamicin treatment did not alter XPC or p16^INK4A^ levels in control fibroblasts (Fig.4E). In line with this, patient fibroblasts treated with gentamicin and transfected with the CDKN2A-luciferase construct produced significantly higher luciferase signals compared to untreated controls (Fig.4F). These results confirmed that rescuing XPC expression by PTC read-through could rescue *CDKN2A* dysregulation, offering a potential treatment and/or melanoma prevention strategy for patients with similar conditions.

## DISCUSSION

We identified a novel role for XPC beyond its established function in DNA repair, demonstrating its critical role in regulating *CDKN2A* expression, particularly in melanocytes. While XPC deficiency typically manifests as increased UV sensitivity and a predisposition to skin cancers(33), our results highlight a novel mechanism of melanomagenesis independent of classic NER defects, with a potential reversal with gentamicin treatment, a clinically available medication. Our study integrates clinical research with basic science investigations to elucidate XPC function and uses this understanding to propose avenues for therapeutic intervention via PTC read-through.

Our identification of two XPC variants in the patients, one previously reported to disrupt splicing(11,12) and another novel splice site variant, provided a unique opportunity to investigate XPC’s DNA repair-independent function. Notably, while both parents carried one variant each and exhibited NER deficiency, only the patients who inherited both variants developed multiple melanomas. Interestingly, several case reports demonstrate similar clinical findings in patients with *XPC* missense variants or late loss-of-function variants, indicating that the phenomenon described herein likely affects other patients(34,35). The clinical findings were then combined with the development of an Xpc^var/var^ mouse model, which mirrors the human splicing variant, allowing for an in vivo investigation. Surprisingly, these mice did not exhibit overt phenotypic abnormalities or increased cancer susceptibility. However, single-cell RNA-seq analysis revealed a striking absence of *Cdkn2a-*expressing melanocytes in the Xpc^var/var^ mice. Interestingly, only 11 of the 25 cell clusters in the mouse tail skin showed *Cdkn2a* expression; furthermore, of the 11 cell clusters that expressed *Cdkn2a*, three showed comparable *Cdkn2a* expression between the WT and the Xpc^var/var^ mice, and six clusters showed increased *Cdkn2a* levels in the WT compared to that in the Xpc^var/var^ mice. Notably, two clusters, IFE Suprabasal-Sebaceous gland and IFE Suprabasal Upper HF 1, showed higher *Cdkn2a* expression in the Xpc^var/var^ mice (Fig.S3B). This strongly suggests a cell-type-specific role for XPC in regulating *CDKN2A* expression, which is potentially crucial for melanoma suppression. Notably, mice with *Cdkn2a* loss do not spontaneously develop melanomas(36,37), indicating that additional factors are likely necessary for melanoma initiation in a murine model.

Despite identifying a novel XPC exon 13 splicing variant (c.2420+5G>A) in patients with multiple melanomas, which led to the discovery of a novel XPC function, we were, unfortunately, unable to detect the predicted truncated protein by western blot or proteomic analysis. RNA-sequencing analyses suggested the presence of multiple XPC transcripts, including those retaining intron 13 (Fig.1C,S1B). While both patients and their parents exhibited significant NER deficiency compared to healthy controls, the patients who inherited two XPC variants demonstrated only subtle reductions in NER capacity compared to their unaffected parents. This suggests that while the XPC variants impaired NER function, the residual XPC protein levels in the patients were sufficient to maintain basic NER activity, which protected them from an overt XP phenotype. Alternatively, the altered XPC protein, potentially truncated or incorporating intronic sequences, might retain NER function while lacking transcriptional activity. While SDS-PAGE and western blot analyses did not reveal significant molecular weight differences, a more detailed mass spectrometry analysis is required to characterize the precise structural alterations of the altered XPC protein. Importantly, this finding highlights the possibility of melanomagenesis in individuals with XPC variants, even with preserved basal levels of NER function.

Our mechanistic studies revealed a novel role for XPC in melanomagenesis by demonstrating its direct involvement in *CDKN2A* regulation. XPC was shown to bind to the *CDKN2A* promoter and is essential for its transcriptional activation, establishing a critical link between two key genes involved in melanoma susceptibility. This finding, coupled with previous research highlighting the importance of functional gene networks in melanoma development(38), underscores the significance of these interconnected pathways in melanoma pathogenesis. These results suggest novel avenues for chemoprevention and drug development targeting XPC-mediated *CDKN2A* regulation. Moreover, rescuing XPC expression in patient fibroblasts through gentamicin-mediated read-through restored *CDKN2A* expression, suggesting a potential therapeutic avenue for patients with similar XPC variants. While systemic gentamicin therapy for melanoma chemoprevention would likely be limited by long-term antibiotic use, stewardship, and possible side effects, our finding offers two future options, including topical gentamicin therapy to higher-risk body sites and a theoretical basis for further drug development.

Previous studies have demonstrated that XPC interacts with RNA polymerase II, contributing to transcription initiation(26,39–41). Moreover, XPC has been reported to modulate chromatin structure by influencing DNA methylation at gene promoters(42,43). While ChIP-sequencing studies have identified genes positively and negatively regulated by XPC, these studies often involve specific cellular contexts or treatments, such as ATRA treatment, potentially limiting their applicability to physiological conditions(26). This highlights the need for comprehensive ChIP-sequencing analyses in various cell types under physiological conditions to gain a more complete understanding of XPC’s transcriptional regulatory landscape. The precise mechanisms by which XPC selectively regulates distinct genes remain elusive. While our work identifies XPC’s transcriptional regulatory role on *CDKN2A* in melanocytes and its importance in melanomagenesis, XPC loss-of-function is relevant in other cell types, leading to other cancer types. While this may be secondary to the loss of DNA repair, XPC likely regulates the transcription of other genes in these cell types, suggesting the need for future comprehensive evaluations of this process. Further, instead of recognizing specific DNA sequences, XPC may recognize broader distortions in DNA conformation, a characteristic also observed in its DNA repair function(29), potentially recruiting other factors to the transcriptional machinery. However, how XPC distinguishes between different promoters and exerts differential effects on gene expression remains unclear. It is hypothesized that XPC’s specificity might be mediated through interactions with distinct sets of protein partners. Future research should prioritize the identification of these XPC-interacting proteins to elucidate the molecular mechanisms underlying their gene-specific regulatory activity.

This study significantly advances our understanding of XPC function in melanoma development. However, further studies generating and analyzing Xpc^var/var^ mice crossed with *Cdkn2a*-deficient mice will be crucial to definitively establish the role of *CDKN2A* in the observed human phenotype. A previous study showed that XPC loss, in conjunction with Ink4a-Arf inactivation, drives melanomagenesis; however, Xpc^-/-^Ink4a-Arf^+/+^ mice failed to develop tumors(37), suggesting that XPC is not the sole transcriptional regulator of *CDKN2A*. Animal models with tissue-specific XPC deficiency are required for a more precise investigation of XPC’s role in cell-type-specific transcription functions.

Finally, the findings of this study have significant implications for our understanding of melanomagenesis and shed light on the transcriptional regulation of CDKN2A, which is largely unexplored. The identification of XPC as a crucial regulator of *CDKN2A* expression in melanocytes expands our knowledge of the molecular mechanisms underlying tumor suppression. This discovery opens new avenues for research, including investigating the precise mechanisms by which XPC interacts with the *CDKN2A* promoter and the factors that modulate this interaction. Furthermore, exploring the potential therapeutic implications of targeting XPC-mediated *CDKN2A* regulation in melanoma warrants further investigation.

In conclusion, this study reveals a novel role for XPC as a crucial regulator of *CDKN2A* expression, particularly in melanocytes. These findings have significant implications for our understanding of melanomagenesis and suggest potential therapeutic targets for patients with XPC and *CDKN2A*-related disorders.

## METHODS

### Experimental model details

#### Patients

Two patients who presented with multiple melanomas at University Hospitals, Cleveland Medical Center, and their parents were enrolled in the "Characterization of the genetic basis for families presenting with multiple melanomas" (Case 3614) study. Subjects had to meet one of the following inclusion criteria: (i) Persons 18 years or older with a confirmed diagnosis of multiple (≥3 biopsy-confirmed) primary melanomas or (ii) Persons 18 years or older who are unaffected first-degree relatives of a person diagnosed with multiple melanomas. The exclusion criteria were as follows: (i) Persons 17 years or younger with a confirmed diagnosis of multiple melanomas; (ii) Persons 17 years or younger who are unaffected first, second, or third-degree relatives of a person diagnosed with multiple melanomas; (iii) Persons who are relatives of a person with a confirmed diagnosis of multiple melanomas and are pregnant at the time of recruitment; or (iv) Non-English speaking individuals. Blood was collected for whole-exome and/or whole-genome sequencing. Punch biopsies were obtained of normal, unaffected skin to obtain fibroblasts for functional studies. All study participants provided a signed informed consent. This study was approved by the Institutional Review Board at University Hospitals Cleveland Medical Center and through Reliant Review at Cleveland Clinic.

#### Mouse model development and validation

A mouse strain carrying the XPC splicing variant, representative of the human XPC variant identified in the patient family, was developed by Cyagen Biosciences Inc. The gRNA to mouse *Xpc* gene, the donor oligo containing c.2399+5G to A mutation, and Cas9 were co-injected into fertilized C57BL6 mouse eggs to generate targeted knock-in offspring. F0 founder animals, identified by PCR followed by sequence analysis, were bred to WT mice to test germline transmission and F1 animal generation. The gRNA target sequence used was (matching reverse strand of gene): 5′ GACCCCTCACGCACACTGGATGG 3′. Additional information on the mouse strain, the breeding strategy, and the genotyping strategy are provided in Fig.S2. The mice carrying the *Xpc* variant did not show any phenotypic differences from the WT mice and did not develop any adverse conditions with aging.

#### Animal experiments

The animal experiment protocol was approved by the Institutional Animal Care and Use Committee of the Cleveland Clinic and was in accordance with the Animal Welfare Act (AWA) and Public Health Service (PHS) Policy of the United States of America. Wild-type (WT) C57BL/6 (RRID: IMSR_JAX:000664) mice purchased from Jackson Laboratories, Wild-type C57BL6 mice (WT) C57BL6 mice heterozygous for the *Xpc* variant (*Xpc^WT/var^)*, and C57BL6 mice homozygous for the *Xpc* variant (*Xpc^var/var^)* were housed in groups of not more than five, in plastic cages with stainless-steel grid tops in the animal care facility of the Cleveland Clinic Lerner Research Center with water and food supplied *ad libitum* in a 12 h light/dark cycle. Mice were euthanized, and their tails were collected and processed as described previously(18)for downstream single-cell RNA-sequencing analysis.

#### Cell lines

HEK-293T and HEK-293 cells and human melanoma cell lines, WM-164, and SK-MEL-28, were obtained from the American Type Culture (ATCC) and were maintained in high-glucose Dulbecco’s modified Eagle’s medium (DMEM; Invitrogen, #11995073), supplemented with 10% fetal bovine serum (FBS; Invitrogen, #26140079) and antibiotic–antimycotic mixture (Invitrogen, 15240096) (100 units/ml) at 37 °C and 5% CO_2_.

#### Patient-derived fibroblasts

Skin punch biopsies were obtained from patients and immediately placed in culture media at 4 °C for temporary storage. Biopsies were sliced into small pieces using a sterile scalpel and plated in 6-well plates. A sterile glass coverslip was placed on each biopsy, and 2.5 ml of culture media was added. Plates were incubated at 37 °C with 5% CO_2_ for 5 days, without disturbances, after which the media was changed every 3 days. Once confluent, fibroblasts were passaged and maintained in high-glucose DMEM (Thermo Scientific, 11960044) supplemented with 10% FBS, 1X GlutaMAX Supplement (Gibco, #35050061), and 1X MEM Non-Essential Amino Acids Solution (Gibco, #11140050).

#### XPC knockdown and overexpression

XPC shRNA-carrying lentiviral particles were produced by transient transfection of HEK-293T cells with a VSV-G envelope expressing plasmid, pMD2.G (a gift from Didier Trono; Addgene plasmid #12259; http://n2t.net/addgene:12259; RRID: Addgene_12259), a 2^nd^ generation lentiviral packaging plasmid, pCMVR8.74 (a gift from Didier Trono; Addgene plasmid #22036; http://n2t.net/addgene:22036; RRID: Addgene_22036), and either XPC shRNA TRCN0000307194 (Thermo Fisher; #NM_004628) (targeting the coding sequence (CDS) CCCACTGCCATTGGCTTATAT; Sh1) or TRCN0000296565 (Thermo Fisher, #NM_004628) (targeting the 3′ UTR ATGGAAGCCACCGGGAGATTT; Sh2), using Lipofectamine 3000 (Thermo Fisher, #13778075), following the manufacturer’s protocol. Lentiviral supernatants were collected 24 h and 48 h after transfection and stored at -80 °C until use. HEK-293T, WM-164, and SK-MEL-28 cells were cultured in fresh culture media containing 8 μg/ml polybrene (Santa Cruz, #sc-134220) and infected with the lentiviral supernatant. Stable XPC-KD cells were selected by culturing the cells with 2.5 μg/ml puromycin (Invivogen, #ant-pr-1) for 4 days. The stable cell lines were maintained in complete DMEM, and cell stocks were prepared and stored in liquid nitrogen. For transient knockdown, the cells were transfected with the plasmid constructs using Lipofectamine 3000, following the manufacturer’s instructions.

Human *XPC* mRNA was amplified from Human Epidermal Melanocyte-adult Total RNA (ScienCell Research Laboratories, Carlsbad, CA. # 2235) using KAPA HiFi HotStart ReadyMix PCR Kit (Roche, #07958927001) and the following primers: Forward primer: AGCAACATGGCTCGGAAA, Reverse Primer: CTGCCTCAGTTTGCCTTCT. XPC mRNA was cloned into the StrataClone PCR cloning vector (Agilent, #240207) and, after sequence confirmation, was moved to pcDNA3.1/Hygro(+). Cells were transfected with the plasmid construct using Lipofectamine 3000, following the manufacturer’s instructions.

### RT-qPCR

Total RNA was isolated from cells using the RNeasy Kit (Qiagen, #74104) following the manufacturer’s instructions. RNA (0.5 µg) was reverse transcribed (RT) to cDNA using random hexamers and the SuperScript III First-Strand Synthesis System (Invitrogen, #18080-051). Quantitative RT-PCR (RT-qPCR) for *XPC* and *CDKN2A* were performed using the following primer sets. *XPC*: Forward: CTGCCATCCTTGGGTATTGT; Reverse: CCTCACCACTCTTGCTTTCT; *CDKN2A*: Forward: ATATGCCTTCCCCCACTACC; Reverse: CACATGAATGTGCGCTTAGG. A five-point, five-fold dilution series was used for primer validation. The absence of non-specific amplification was confirmed by a single peak in the melt-curve analysis; the amplicon size and the absence of amplification in the no template control were confirmed by agarose gel analysis. qPCR was performed in triplicate on the StepOne Plus (Applied Biosystems CA, USA) using the PowerUP SYBR Green Master Mix (Applied Biosystems, #A25741). Cycling conditions were: 50 °C for 3 min, 95 °C for 5 min, followed by 40 cycles of 95 °C for 15 s and 60 °C for 30 s. The relative mRNA levels were calculated using the 2^−ΔΔCT^ method using *GAPDH* as the control.

### Evaluation of DNA repair capacity

Patient-derived fibroblasts and normal human skin fibroblasts OSU-2 were cultured in 60-mm dishes to confluence, maintained in serum-free medium for 12 h, treated with ultraviolet-C (UVC) radiation at 10 J/m^2^, and further cultured in serum-free medium for the desired periods (0, 8, 24, or 32 h). Cells were harvested by trypsinization, and total genomic DNA was isolated and quantitated. The total DNA (20 ng) was loaded on a nitrocellulose membrane, blocked with 5% non-fat milk, incubated with anti-cyclobutane pyrimidine dimers (CPD) antibody (TDM2, Cosmo Bio LTD, #CAC-NM-DND-001, 1:2000) at 4 °C overnight, washed three times with TBST, incubated with goat-anti-mouse IgG-HRP (Promega, #W402B, 1:5000) at room temperature for 1 h, and subsequently washed three times with TBST. A chemiluminescence substrate was added to the membrane, and CPD bands were detected by film exposure. The film was scanned, and the intensity of each band was quantitated using ImageJ. The relative band intensity was calculated with 0 h time point as 100%.

### Patient RNA-sequencing and analysis

Total RNA was extracted using the QIAamp RNA Blood Mini Kit (Qiagen, # 52304) and quantified using a Qubit Fluorometer (Invitrogen). RNA quality was assessed using the Agilent 2100 Bioanalyzer (Agilent) using a cut-off of RIN > 7.0 to select specimens for further analysis. Library preparation was completed using the Illumina TruSeq Stranded Total RNA kit with Ribo-Zero Gold (Illumina, #20020612) for rRNA removal. This protocol starts by using the Ribo-Zero kit to remove rRNA from 150 ng of total RNA using a hybridization/bead capture procedure that selectively binds rRNA species using biotinylated capture probes. The resulting purified mRNA was used as input for the Illumina TruSeq kit, in which libraries are tagged with unique adapter indexes. Final libraries were validated on the Agilent 2100 Bioanalyzer, quantified using qPCR, and pooled at equimolar ratios. Pooled libraries were diluted, denatured, and loaded onto the Illumina HiSeq 2500 system, following the Illumina User Guide for a paired-end run.

RNA-seq reads were aligned to the reference genome (GRCh38) and transcriptome (gencode.v22) using aligner STAR (2.5.2b), which maps the RNA-seq reads to genomic regions and detect splice junctions by identifying reads that span exon-exon boundaries. The alignment process identifies splice junction reads and computes read coverage for each exon. The alignment BAM files were visualized in Integrated Genomics Viewer (IGV) as Sashimi plots to extract splice junction reads and read coverage. Represented as horizontal bars or curves, showing read density across each exon. Higher coverage indicates higher expression of the exon. Curved lines connecting exons, with their thickness proportional to the number of splice junction reads supporting that specific splice event. This visually represents the frequency of specific splicing events.

### Single-cell RNA-sequencing

Tail skin samples from three WT and three *Xpc^var/var^ mice* were collected and processed as described previously(18) to obtain highly viable single-cell suspensions for single-cell RNA-seq analysis. Single-cell capturing and library preparation were performed using the 10x Genomics Chromium Next GEM Single-Cell 3 Reagent Kits v3.1 (#CG000204) following the manufacturer’s protocol. The library was sequenced using the Illumina NovaSeq platform. Sequence data were analyzed, and optimal cell clusters were identified using the shared nearest neighbor (SNN) method(17), as described previously(18). Data visualization and subsequent analyses were performed using BBrowser (BioTuring).

### Flow cytometry

Cells fixed and permeabilized using the Inside Stain Kit (Miltenyi Biotec, #130-090-477), following the manufacturer’s protocol, were incubated with DyLight 594-conjugated human XPC antibody (Boster Biological Technology, #A00473-1-Dyl594) and Alexa Fluor 488-conjugate CDKN2A/p16INK4a antibody (Bioss Antibodies, #bs-4592R-A488) for 10 min at room temperature in the dark, after which, fluorescence data from triplicate samples were acquired on a BD Fortessa (BD Biosciences) and analyzed using FlowJo v10.10.

### Cell growth and tumorigenicity

To determine cell growth, 15,000 cells in 100 μl media were plated in each well of a 96-well plate and incubated at 37 °C. At specific time points after seeding, 10 μl WST-8 solution (Colorimetric Cell Viability Kit 1, PromoKine, #PK-CA705-CK04) was added to each well. The plates were incubated at 37 °C for 45 min, and the absorbance at 450 nm was read using a Varioskan LUX multimode microplate reader (Thermo Fisher). The average value from triplicates was used to determine cell viability.

In vitro, tumorigenicity was evaluated as described previously(25). Cells were suspended in 0.35% agarose-DMEM and layered on 0.5% agar-DMEM in 6-well plates. Complete media (1 ml) was layered over the agarose layer, and the cells were cultured in a humidified incubator at 37 °C and 5% CO_2_. Fresh media was added to the wells every 3 days. After 3 weeks, the gels were stained with crystal violet (Sigma-Aldrich, #HT90132-1L), and the plates were observed under a wide field microscope (Leica DM5500B, Leica Microsystems, GmbH, Wetzlar, Germany equipped with a Leica DFC425C color camera (Leica Microsystems, GmbH, Wetzlar, Germany) and the LASX software. The number of colonies in each well was counted using Image-Pro Plus (Media Cybernetics, Inc., Rockville, MD). The average from triplicate wells was used for calculations, and control wells with no cells were used to eliminate the background.

### Western blot

Total cell and tissue lysates were prepared using M-per Mammalian Protein Extraction Reagent (Thermo Fisher, #78501) containing protease and phosphatase inhibitors (Halt Protease and Phosphatase Inhibitor Cocktail (100X) (Thermo Fisher, #87785)). Equal amounts of proteins were resolved by SDS-PAGE, transferred to nitrocellulose membranes (Bio-Rad, #1620115), blocked with 5% milk or EveryBlot Blocking Buffer (Bio-Rad, #12010020) for 1 h, and incubated with primary antibodies overnight at 4 °C, after which they were washed and incubated with corresponding HRP-conjugated secondary antibodies. The primary and secondary antibodies used in this study were XPC (D1M5Y) Rabbit mAb (Cell Signaling Technology, #14768), CDKN2A/p16 Antibody (JC8) (Sanga Cruz, #sc-56330), GAPDH (14C10) Rabbit mAb (Cell Signaling Technology, #2118), HRP-conjugated Anti-Rabbit IgG (Promega, #W401B), and, HRP-conjugated Anti-Mouse IgG (Promega, #W402B). The protein bands were developed using the ECL Western Blotting Substrate (Bio-Rad, Cat. #32106) and visualized using a ChemiDoc imager (Bio-Rad).

### Chromosome immune precipitation (ChIP)

Chromosome immune precipitation was carried out using the SimpleChIP Plus Enzymatic Chromatin IP Kit (Magnetic Beads) (Cell Signaling Technology, #9005), following the manufacturer’s protocol. Briefly, HEK-293T cells were crosslinked with formaldehyde to preserve protein-DNA complexes. The nuclei were isolated, and the chromatin was enzymatically fragmented. XPC antibody (D-10) (Santa Cruz, #sc-74410) was used to capture XPC-DNA complexes, which were then isolated using magnetic beads. An anti-GFP antibody (GFP (B2); Santa Cruz, #sc-9996) was used as the negative control. The protein-DNA complexes were eluted from the beads, and the crosslinks were reversed to release the DNA. RT-qPCR for *CDKN2A* promoter sequences was performed using the following primers and the SimpleChIP Universal qPCR Master Mix (Cell Signaling Technology, #8898). The following primers against the *CDKN2A* promoter regions were used: Primer set 1: Spans chr9:21,994,139-21,994,39: Forward: GGTCCCAGTCTGCAGTTAAG; Reverse: AAGAACCTGCGCACCAT. Primer set 2: Spans chr9:21,994,139-21,994,39: Forward: CTCGTGCTGATGCTACTGA; Reverse: TTTCGAGGGCCTTTCCTAC. Primer set 3: Spans chr9:21,974,686-21,974,809: Forward: AGCAGCATGGAGCCTTC; Reverse: GCCTCCGACCGTAACTATTC. Primer set 4: Spans chr9:21,974,686-21,974,809: Forward: ACAGAGTGAACGCACTCAAA; Reverse: GTTGGCAAGGAAGGAGGAC. The absence of non-specific amplification was confirmed by a single peak in the melt-curve analysis. The cycling conditions were as follows: 95 °C for 3 min, followed by 40 cycles of 95 °C for 15 s and 60 °C for 60 s. IP efficiency was calculated using the Percent Input Method.

### Luciferase assay

The LightSwitch Promoter Reporter system was used to determine promoter binding. Cells seeded in white 96-well tissue culture plates were transfected with either the *CDKN2A* promoter construct (SwitchGear Genomics, #32001) or the Random promoter control construct (SwitchGear Genomics, #320070) using the FuGENE HD Transfection Reagent (Promega, #E2311), following the manufacturer’s protocol. After 48 h, the luciferase reporter signal was determined using the LightSwitch Luciferase Assy Kit (SwitchGear Genomics, #32031) and a Varioskan LUX multimode microplate reader (Thermo Fisher), following the manufacturer’s protocol.

### Statistical analysis

All statistical analyses were performed using GraphPad Prism Ver 10.1.12 (GraphPad Software, La Jolla, CA, USA), R (version 4.0.3), and RStudio (version 1.3.1093). The specific tests used for each analysis are described in the Figure legends. A p-value <0.05 was considered statistically significant.

## ACKNOWLEDGMENT

We thank Harley and Rochelle Gross and the Gross Family for funding this study and their philanthropic support.

This research was supported by the Genomics Core Facility of the Cleveland Clinic Lerner Research Center and the Case Western Reserve University School of Medicine.

Plasmids pMD2.G (RRID: Addgene_12259) and pCMVR8.74 (RRID: Addgene_22036) were gifts from Didier Trono.

## ETHICS STATEMENT

This study was conducted in accordance with the ethical principles outlined in the Declaration of Helsinki and was approved by the Institutional Review Board (IRB) at the University Hospitals Cleveland Medical Center and through Reliant Review at Cleveland Clinic (Case 3614). All participants provided written informed consent prior to participating. All animal experiment protocols were approved by the Institutional Animal Care and Use Committee of the Cleveland Clinic and were in accordance with the Animal Welfare Act (AWA) and Public Health Service (PHS) Policy of the United States of America.

## DATA AVAILABILITY STATEMENT

The RNA-seq and the single-cell RNA-seq data will be submitted to a public repository, and the accession numbers will be provided once the manuscript is accepted for publication.

