## Supplementary material for "XPC loss-of-function triggers melanomagenesis through *CDKN2A* downregulation": S1

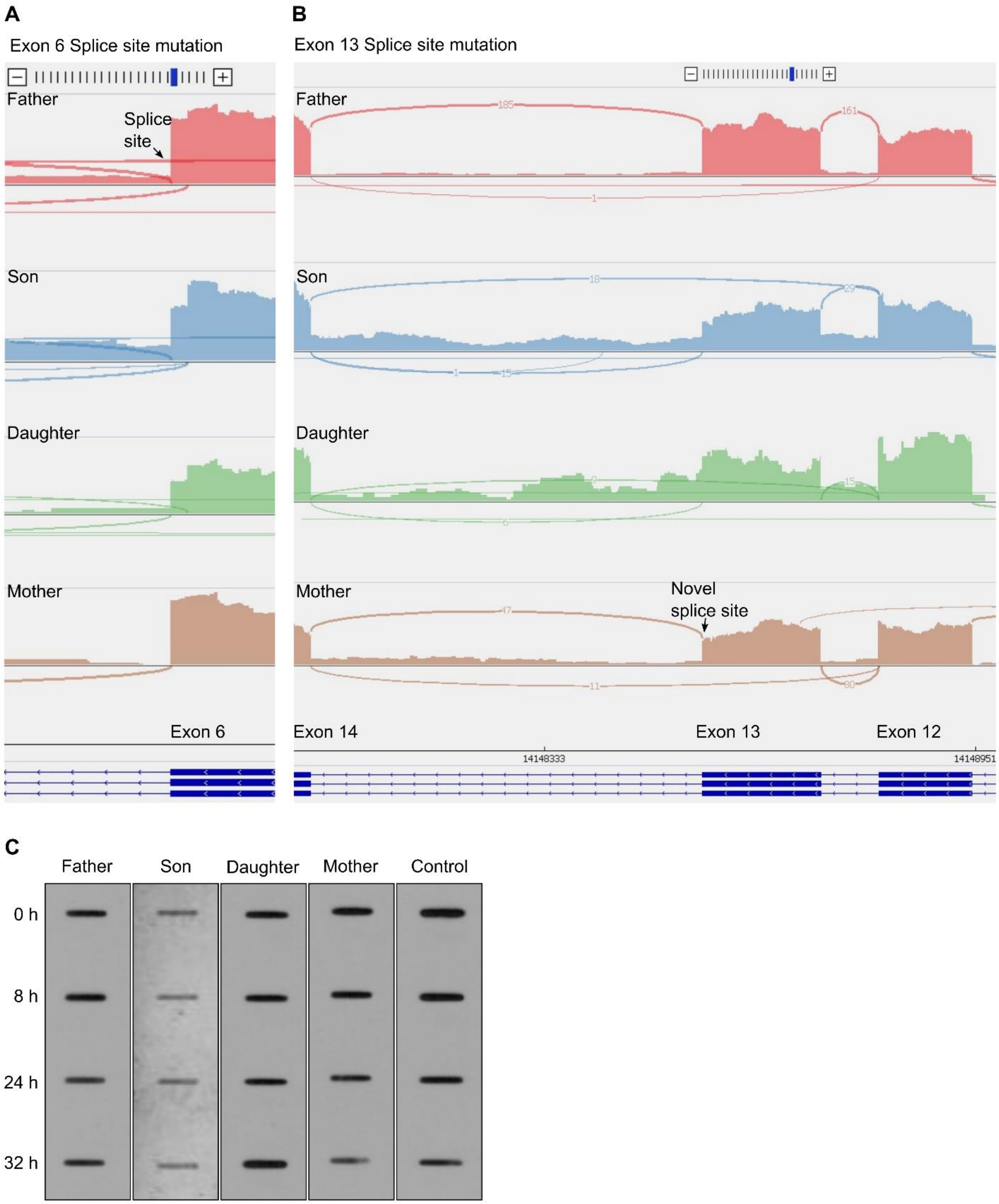


**Figure S1. (A, B)** Zoomed-out sashimi plots of the possible alternative splicing products of XPC (**A**) for variant C779+1G>T, the exon 6 splice site variant inherited from the father, and (**B**) C240+5G>A, the novel exon 13 splice site variant identified in this study and inherited from the mother. (**C**) Representative gel image showing the cyclobutane pyrimidine dimers (CPD) blotted using an anti-CPD antibody in lysates of fibroblasts derived from the patients and their parents and normal human skin fibroblasts (OSU-2, Control) at different time points after ultraviolet-C (UVC) irradiation.
