## Supplementary material for "XPC loss-of-function triggers melanomagenesis through *CDKN2A* downregulation": S2

**A**

**B**

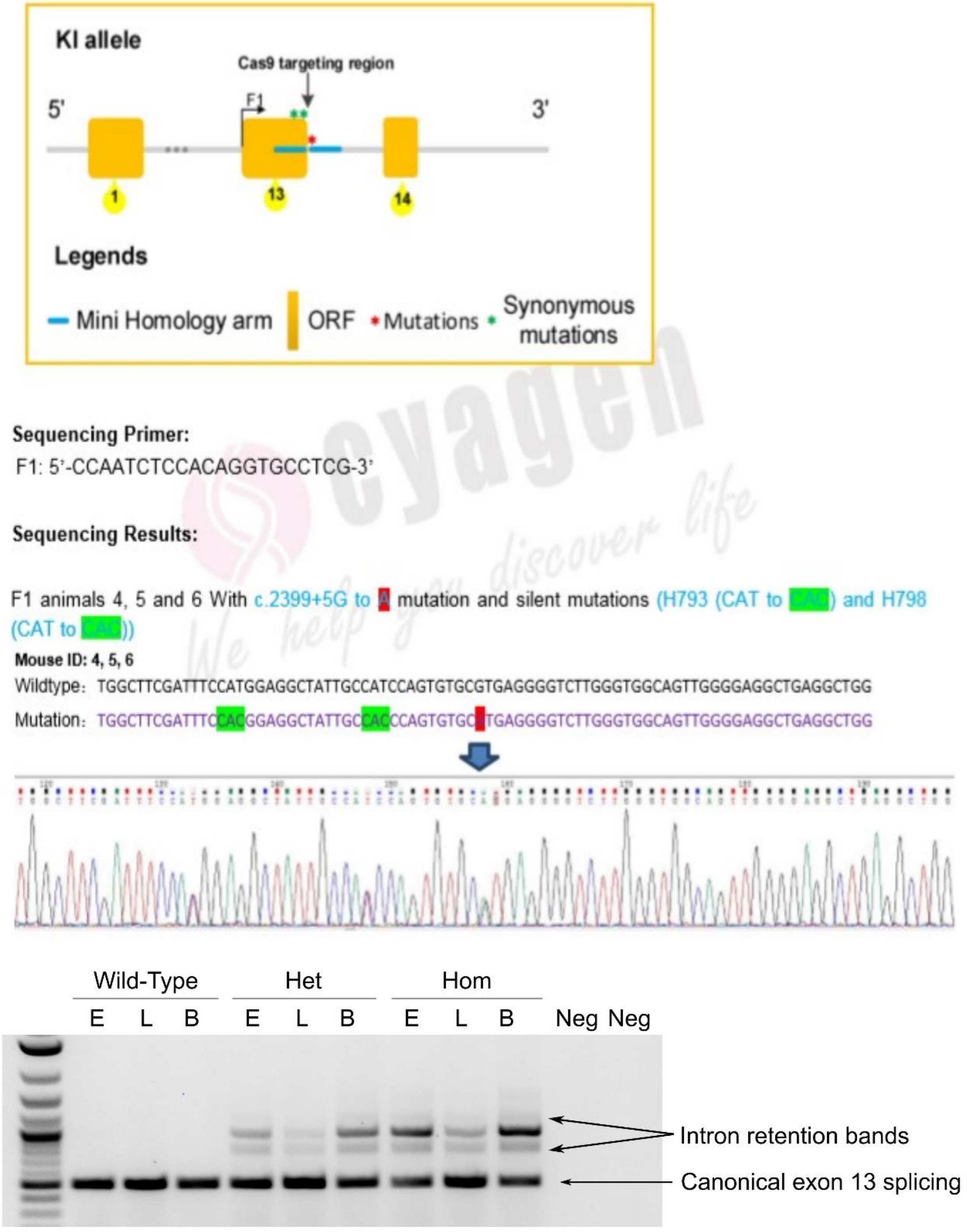


**Figure S2**. Development of a mouse model carrying the *Xpc* splicing variant, representative of the human *XPC* variant identified in the patient family. (**A**) Scheme showing the sequencing strategy and sequencing confirmation of the mouse model. (**B**) PCR determination of *Xpc* in different tissues (E, Ear; L, Lung; and B, Brain) of the wildtype, heterozygous mutant (Het), and homozygous mutant (Hom) mice carrying the *Xpc* variant.
