## Supplementary material for "XPC loss-of-function triggers melanomagenesis through *CDKN2A* downregulation": S3

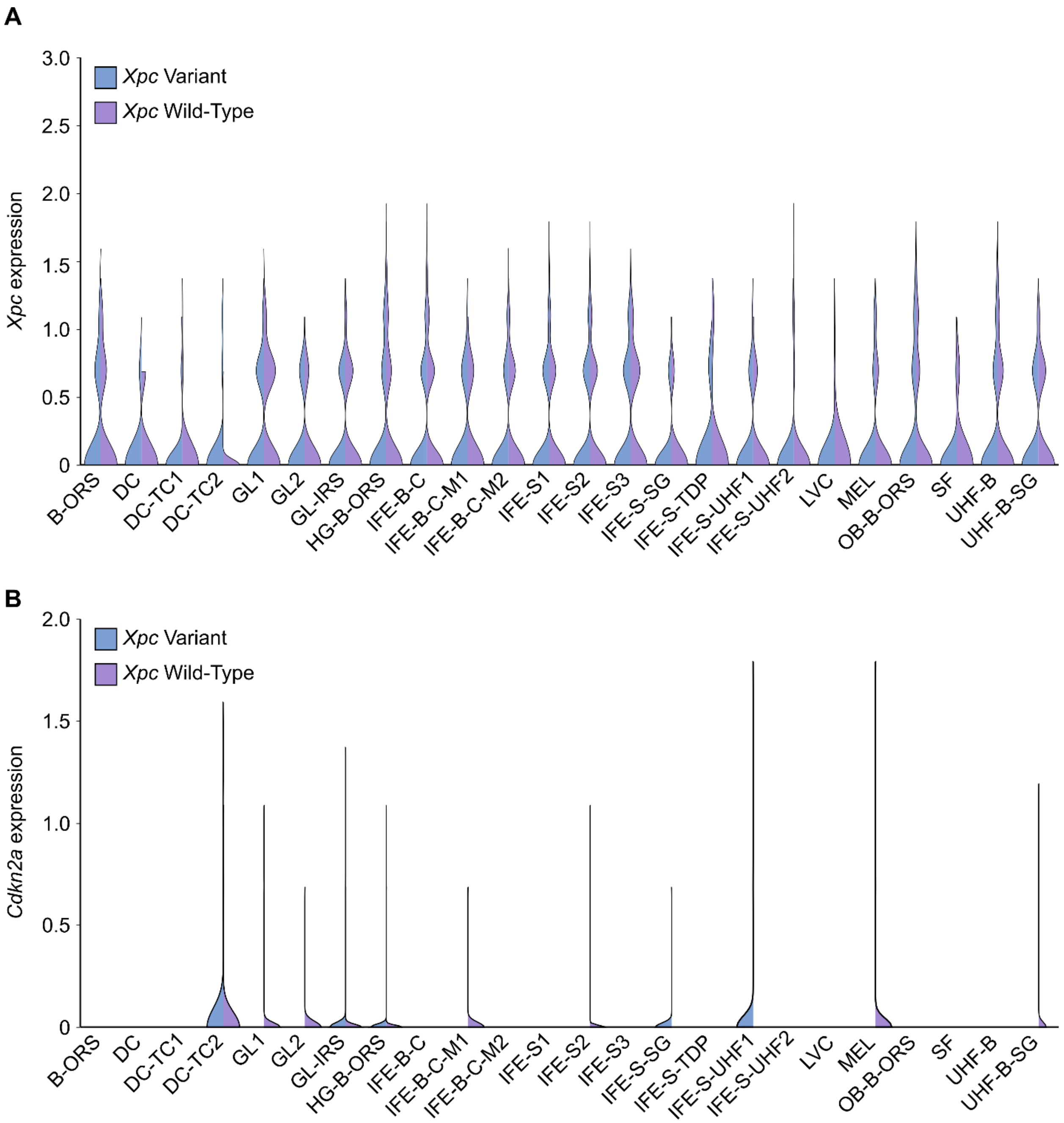


**Figure S3**. Violin plot showing the expression of (**A**) *Xpc* and (**B**) *Cdkn2a* in different cell clusters of the wild-type and Xpc$^{var/var}$ mice.
